# BioWiC: An Evaluation Benchmark for Biomedical Concept Representation

**DOI:** 10.1101/2023.11.08.566170

**Authors:** Hossein Rouhizadeh, Irina Nikishina, Anthony Yazdani, Alban Bornet, Boya Zhang, Julien Ehrsam, Christophe Gaudet-Blavignac, Nona Naderi, Douglas Teodoro

## Abstract

Due to the complexity of the biomedical domain, the ability to capture semantically meaningful representations of terms in context is a long-standing challenge. Despite important progress in the past years, no evaluation benchmark has been developed to evaluate how well language models represent biomedical concepts according to their corresponding context. Inspired by the Word-in-Context (WiC) benchmark, in which word sense disambiguation is reformulated as a binary classification task, we propose a novel dataset, BioWiC, to evaluate the ability of language models to encode biomedical terms in context. We evaluate BioWiC both intrinsically and extrinsically and show that it could be used as a reliable benchmark for evaluating context-dependent embeddings in biomedical corpora. In addition, we conduct several experiments using a variety of discriminative and generative large language models to establish robust baselines that can serve as a foundation for future research.

## 1. Background & Summary

Biomedical corpora, such as scientific articles and patient reports, contain a wealth of knowledge and information that can be used to enable high-quality research. However, the extraction of knowledge from these free-text sources is a challenging task as it requires the ability to understand the meaning of natural language and the idiosyncrasies of the biomedical domain but also due to the volume of the data^1^. Biomedical natural language processing (NLP) techniques have been used to analyze information from free-text sources at scale, enabling the extraction and synthesis of biomedical information, and transforming unstructured data into a structured format^2,3^.

Compared to general corpora, NLP models face three main challenges for semantic representation of biomedical data^4–7^. First, the number of biomedical entities is extremely high. For example, the SNOMED-CT ontology^8^ defines more than 300’000 medical concepts while the UniProt Knowledgebase (UniProtKB)^9^ contains more than 550’000 curated proteins. Combined, the number of concepts described in these two knowledge organization systems is higher than the number of terms defined in dictionaries for many natural languages. Second, biomedical concepts have many synonyms and alternative expressions for the same concept. For example, in Figure 1 the concept *“C0007134”* defined in the Unified Medical Language System (UMLS) thesaurus can be represented with at least four terms: *“Renal Cell Carcinoma”, “RCC”, “Nephroid Carcinoma”*, and *“Adenocarcinoma”*. Third, biomedical corpora are notorious for their overabundance of abbreviations and acronyms^10^. These abbreviations and acronyms are often polysemous, e.g., the acronym *“RCC”* in Figure 1 belongs to two concepts – *“C2826323”* and *“C0007134”* – making their semantic representation even more challenging.

**Figure 1.**
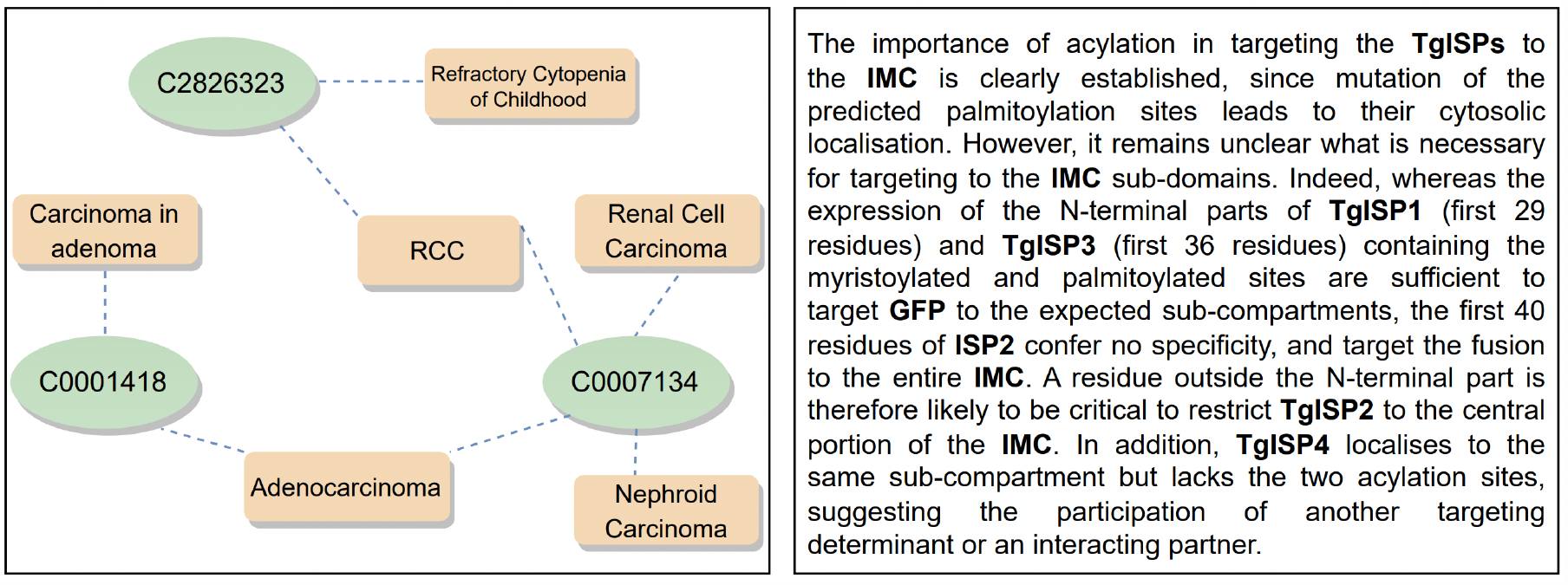
Illustration of concept ambiguity in the biomedical domain. Left: Example of the UMLS 2021AB data structure, where one term refers to different concepts as well and one concept may be represented with different mentions. Right: Example of a paragraph with numerous polysemous acronyms and abbreviations from a biomedical journal ^11^. Acronyms and abbreviations are highlighted in bold.

Word sense disambiguation (WSD)^12^ and entity linking^13^ are two NLP tasks trying to address the issue of semantic representation in the biomedical field. Given a word in context, the objective of WSD is to associate the word with its correct meaning in a sense inventory^14,15^. For example, in the sentence *“The patient has been suffering from a cold*.*”*, the sense for the word *cold* should be associated with its *medical* meaning as opposed to *temperature* or *literature* (i.e., James Bond novel by John Gardner) meanings. On the other hand, entity linking aims to connect terms mentioned in a text with corresponding concepts in a knowledge organization system^16,17^. For instance, the abbreviation *“CA”* in biomedical contexts can stand for either *“calcium”*, an essential mineral in the human body, or *“cancer”*, a group of diseases characterized by abnormal cell growth. An ideal entity linking system would employ contextual cues to correctly map *“CA”* to its standardized form in a chosen knowledge base, e.g., UMLS. This proper alignment assists in reducing ambiguity, enhancing the understanding of biomedical corpora^18,19^.

Recently, the Word-in-Context (WiC) benchmark^20^ presented a novel perspective on WSD. WiC formulates WSD as a binary classification task, where a polysemous word appears in two different sentences, and the task is to infer whether the word holds the same meaning or not. WiC has been integrated as a component of SuperGLUE^21^, a comprehensive evaluation framework designed to assess the performance of natural language understanding systems. XL-WiC^22^ and TempoWiC^23^ are two recent extensions of WiC adapting it to 12 different languages and targeting the detection of meaning shifts in Twitter, respectively. Despite significant progress both in WSD and entity linking tasks in the biomedical domain^18,24–26^, there exists no benchmark that specifically targets the semantic representation of biomedical terms in context.

To bridge this gap, we present the BioWiC benchmark, a novel dataset that provides high-quality annotations for the evaluation of contextualized term representations in the biomedical domain. Inspired by the WiC^20^, we formulate BioWiC as a binary classification task, whose aim is to identify whether two target terms in their respective contexts have the same meaning. In addition to its focus on biomedical concepts, BioWiC differs from WiC in several ways. First, in contrast to WiC which focuses on single token words, as targets, BioWiC allows for terms that can be single words, phrases, or multiword expressions. Second, BioWiC terms may be represented not only by the same terms in different contexts but also by different term forms referring to the same concept (or not). The dataset is named “BioWiC”, reflecting its design for the biomedical domain while showcasing its relation to the WiC task.

## 2. Methods

In this section, we present BioWiC – the first benchmark dataset for evaluating in-context biomedical concept representations. First, we explain the resources we used to create the corpus and the pre-processing steps. We then provide an overview of the methodology used to create the dataset and discuss the processes for instance generation, dataset splitting, and quality assessment.

### 2.1 BioWiC resources

As shown in Table 1,BioWiC instances were built using annotations from the following manually curated biomedical entity linking datasets:

**Table 1.** General statistics of BioWiC resources. The sentence count in each source is determined using the PySBD library^32^, version 0.3.4.

| Dataset | Ontologies | Semantic types | Documents | Sentences | Mentions |
| --- | --- | --- | --- | --- | --- |
| Medmentions | UMLS | 21 UMLS types | 4392 | 44903 | 203'282 |
| BC5CDR | MeSH | Disease, Chemical | 1500 | 11562 | 13'343 |
| NCBI Disease | MeSH, OMIM | Disease | 792 | 3891 | 6'892 |

**MedMentions**^**27**^: this is the largest entity linking dataset in the biomedical domain. It includes 4’392 PubMed abstracts and over 350’000 mentions linked to UMLS. The full MedMentions version covers 128 UMLS semantic types. However, as stated by^27^, the concepts can be either too expansive (e.g., “Group, South Asia”) or cover peripheral and supplementary topics (e.g., “Rural Area, No difference”). Thus, we follow^28,29^ and focus on the officially released subset of MedMentions called ST21pv (21 Semantic Types from Preferred Vocabularies), which contains 203’282 biomedical mentions from 21 UMLS semantic types.

**BC5CDR**^**30**^: introduced in the BioCreative challenge, this dataset comprises 1’500 PubMed abstracts and 13’343 mentions linked to Medical Subject Headings (MeSH) concepts. The dataset covers a wide range of biomedical entities, including 4’409 chemicals, 5’818 diseases, and 3’116 instances of chemical-disease interactions.

**NCBI Disease**^**31**^: developed by the National Center for Biotechnology Information (NCBI), this dataset includes biomedical information derived from 793 PubMed abstracts. It comprises 6’892 disease mentions, each associated with their relevant standardized forms in the MeSH or Online Mendelian Inheritance in Man (OMIM) terminologies.

### 2.2 Data pre-processing

To have homogeneous word-in-context instances from different resources, we unified their format using the following steps:

- **Sentence segmentation**: Each BioWiC instance is composed of a pair of target terms together with their respective sentences. We use the PySBD library^32^, version 0.3.4, to determine sentence boundaries in the initial source texts (i.e., abstracts of publications). We parse documents and keep only sentences that contain mapped mentions.
- **Label unification**: The source datasets of BioWiC map mentions (i.e., terms) to different target knowledge organization resources, i.e., MeSH, OMIM, and UMLS. This results in concept codes, i.e., unique identifiers in the target ontology, that cannot be directly comparable. To address this issue, we used UMLS as the main reference and transferred the concept identifiers from MeSH and OMIM to UMLS using available ontology mappings in UMLS 2021AB. To avoid ambiguity, MeSH or OMIM concepts with multiple mappings in UMLS 2021AB were removed.

### 2.3 BioWiC construction

BioWiC instances follow a similar format to WiC, where each instance involves a pair of biomedical terms (*w*_1_ and *w*_2_) and their corresponding sentences (*s*_1_ and *s*_2_). The task is to classify each instance as *True* if the target terms carry the same meaning across both sentences or *False* if they do not. We represent each instance as a tuple pair *t* = [(*s*_1_,*w*_1_),(*s*_2_,*w*_2_)] : *y* where *w*_1_ and *w*_2_ are the target terms, *s*_1_ and *s*_2_ are the corresponding sentences, and *y* is the associated binary label. Table 2 presents some examples of BioWiC instances. In contrast to WiC, where both target terms of each instance always share the same lemma, BioWiC allows for variations such as abbreviations, synonyms, identical terms, and terms with similar surface forms.

**Table 2.** BioWiC instances, drawn from the test split. The target terms of each instance are in bold.

| Instance group | # | Sentence 1 | Sentence 2 | Label |
| --- | --- | --- | --- | --- |
| Term identity | 1 | ... clinical use of the anthracycline doxorubicin (DOX) is limited by its **cardiotoxic** effects ... | ... mitomycin C (MMC) has been suggested to be **cardiotoxic**, especially when combined with or given ... | T |
|  | 2 | ... a wide range of key concepts and terms of **PE** from clinical and biomedical research ... | ... associated with mortality in COPD patients with low-risk **PE** (adjusted OR 1.11; 95% CI, 1.04-1.66) ... | F |
| Abbreviations | 3 | ... the gene responsible for **FEO** to an interval of less than 5 cM between D18S64 and D18S51... | ... affecting the signal peptide of RANK, cause **familial expansile osteolysis**. | T |
|  | 4 | **Periodontal disease** has risk factors in common with a number of other non-communicable diseases ... | ... allosteric activator of mGlu7 receptors, were thus tested in different rodent models of **PD**. | F |
| Synonyms | 5 | Initial low levels of **IL-10** were associated with an increase in physical disability ... | Assessment of Interleukin-17A, **Interleukin-10** and Transforming Growth ... | T |
|  | 6 | ... variations of heart period (HP), systolic arterial pressure (SAP) and **respiration (R)**. | ... subjects who fail **ventilation** with the C-E technique can be ventilated effectively ... | F |
| Label similarity | 7 | ... deletion of the KIT and PDGFRA genes may account for the **piebald** phenotype in this patient ... | **Piebaldism** in this family thus appears to be the human homologue to dominant white spotting (W) ... | T |
|  | 8 | More **anemic** than non-anemic FDS2 patients died (28.7% versus 8.0%; P<0.001) ... | ... observed were comparable in AZT and PHZ treated mice with similar degrees of **anaemia**. | F |

To evaluate challenging scenarios for semantic representation, such as synonymy, polysemy, and abbreviations, BioWiC is divided into four main groups of instances. Group A (term identity) contains instances where the target terms *w*_1_ and *w*_2_ are identical. In group B (abbreviations), either *w*_1_ or *w*_2_ could represent the abbreviation of the other one. Group C (synonyms), consists of instances where *w*_1_ and *w*_2_ could be synonyms (according to UMLS). Lastly, group D (label similarity) includes instances where *w*_1_ and *w*_2_ share similar surface forms. We employed the following five steps to generate the BioWiC instances:

i. **Sentence collection**: We first gathered all the sentences from the source datasets manually annotated with terms *M*(*W,C*) = {(*w*_1_,*c*_1_),(*w*_2_,*c*_2_),…,(*w*_*n*_,*c*_*n*_)}, where *w* ∈*W* is a term and *c* ∈*C* is a concept defined in UMLS. Then, we created a set *S* = {*s*_1_,…,*s*_*n*_}, where each sentence *s* ∈ *S* has at least one mention *w* ∈*W* linked to *c* ∈*C*.
ii. **Tuple creation**: For each sentence *s* ∈ *S*, we randomly chose one of the annotated mentions *w* and created a set of sentence-term tuples *P* = {(*s*_1_,*w*_1_),(*s*_2_,*w*_2_),…,(*s*_*n*_,*w*_*n*_)}, where for each (*s*_*i*_, *w*_*i*_) ∈ *P, s*_*i*_ includes *w*_*i*_. We then paired the tuples of *P* and created a collection of tuple pairs:

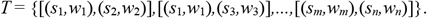
iii. **Instance definition and labeling**: We considered each pair *t* = [(*s*_*i*_,*w*_*i*_),(*s*_*j*_,*w*_*j*_)] ∈ *T* as a potential BioWiC instance, where *w*_*i*_ and *w*_*j*_ serve as target terms and *s*_*i*_ and *s*_*j*_ are their corresponding sentences, respectively. Each instance is labeled as *y* = *True* if the target terms *w*_*i*_ and *w*_*j*_ were linked to the same or synonym UMLS concept, and as *y* = *False* if they were not. We then added the label *y* to each tuple pair to create the dataset of possible BioWiC instances *t =* [(*s*_*i*_,*w*_*i*_),(*s*_*j*_,*w*_*j*_)] : *y*.
iv. **Tuple selection**: We categorized each instance *t* : *y* to one of the main groups of BioWiC. Group A included instances for which *w*_*i*_ and *w*_*j*_ are identical. Group B included instances where *w*_*i*_ is the abbreviated form of *w*_*j*_ or vice-versa. Group C included instances where *w*_*i*_ and *w*_*j*_ could be synonyms. Group D included instances where *w*_*i*_ and *w*_*j*_ are not identical but share similar surface characteristics (see section 2.3.2).
v. **Dataset splitting**: We divided the instances into three parts: training set, development set, and test set, providing a consistent and reliable framework for model training and evaluation (see section 3).

For clarity, in Figure 2 we provide an example of building BioWiC instances for the target term “delivery”. Initially, we preprocess the resource data and extract all sentences in which “delivery” is linked to UMLS. We transform each sentence to the sentence-term tuple (*s*_*i*_,*w*) format where *s*_*i*_ represents a sentence containing the term *w* = “delivery”. Subsequently, we permute all possible combinations of tuples (*s*_*i*_,*w*) identified in the preceding step to generate BioWiC instances *t* = [(*s*_*i*_,*w*),(*s*_*j*_,*w*)], where “delivery” serves as the target term in both sentences. Finally, we classify each instance as *True* when “delivery” is mapped to the same CUI code in both sentences and as *False* when it is not.

**Figure 2.**
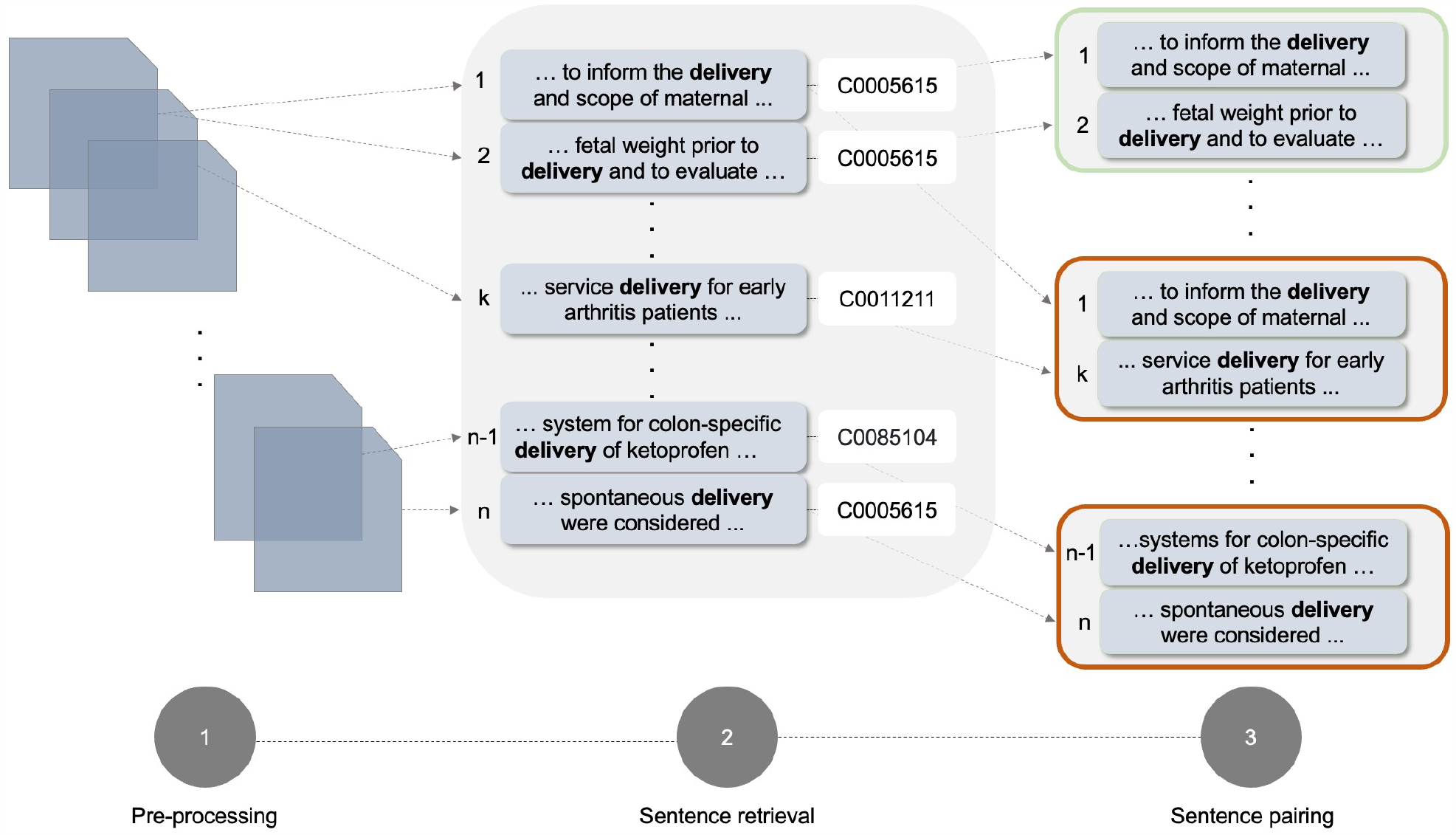
The overall pipeline of the BioWiC construction process. Step 1: Pre-process the source documents to a consistent format. Step 2: Identify and retrieve sentences including the term “delivery” linked to UMLS. Step 3: Pair the retrieved sentences to generate BioWiC instances. In Step 3, the green box shows an example of a BioWiC instance with the same target concept, while the red boxes show examples of different target concepts.

#### 2.3.1 Instance generation

To build the BioWiC instances, we considered two main challenges of biomedical texts: semantic and lexical ambiguities. The presence of *semantically ambiguous terms*, that is, terms that can have multiple meanings in different contexts, is one of the most difficult aspects of biomedical text processing^3^. For instance, the term s*taph* can be used either as a type of *disease* (usually followed by infection) or *bacteria* in other contexts. In addition, one concept can be used in different domains to represent meaning. To assess the capability of language models to provide context-sensitive representations for a term across different contexts, we included a group of instances (group A) in BioWiC in which a target biomedical term appears in two different contexts. Another key challenge in the biomedical domain is that terms can be expressed in various forms or using different *lexical formats*, even if they refer to the same biomedical concepts. To account for this challenge, we developed three other groups of BioWiC instances to measure language models’ ability to use context and produce similar representations for synonym terms with different surface strings. We categorize synonyms into three different groups: *i*) abbreviations, *ii*) synonyms, and *iii*) concepts with similar surface characteristics. Each instance in these groups contains two target terms with different surfaces, each occurring in a different context and the models should identify whether these terms refer to the same biomedical concept or not.

#### 2.3.2 Instance groups

In what follows, we discuss how we created the instances for each group:

A. **Term identity:** To create these instances, we use the tuple pair list, built-in step 3 of the construction pipeline, and consider every pair *t* = [(*s*_*i*_,*w*_*i*_),(*s*_*j*_,*w*_*j*_)] ∈ *T* as an instance of group A if *w*_*i*_ and *w*_*j*_ are identical. We classified each *t* as *True* if both terms were linked to the same UMLS CUI and *False* otherwise. Two instances of this type are shown in Table 2 (examples one and two). In the first example, both target terms refer to the same concept and have the same meaning (i.e., toxicity that impairs or damages the heart, UMLS CUI C0876994). So, the instance label is *True*. In the second instance, however, the target terms are mapped to different CUIs (C0032914 and C0034065), and thus the instance label is *False*.
B. **Abbreviations**: In this group, one of the target terms is the abbreviated form of the other one, e.g., *heart rate* and *hr*. From the tuple pair list, we pick up all the pairs *t* = [(*s*_*i*_, *w*_*i*_),(*s*_*j*_,*w*_*j*_)] ∈ *T* if *w*_*i*_ is the abbreviated form of *w*_*j*_ or vise-versa. To verify this, we generate the abbreviated form of *w*_*i*_ by combining the initial letters from each part obtained after splitting it (e.g., “FEO” is considered as the abbreviation of “familial expansile osteolysis”). Next, we compare whether *w*_*j*_ is the same as the abbreviation of *w*_*i*_. We perform the same procedure for *w*_*j*_ as well. If either of the *w*_*i*_ or *w*_*j*_ is the abbreviation of the other, we categorize the tuple pair into this group. Each tuple pair then is assigned to the label *True* if *w*_*i*_ and *w*_*j*_ are mapped to the same UMLS and *False* otherwise. As shown in example 3 of Table 2, *“FEO”* in sentence 1 is used as the abbreviation of *“familial expansile osteolysis”*. So the instance is labeled as *True*. In example 4, however, the target term *PD* does not have the same meaning as *“Periodontal disease”* and thus the instance is labeled as *False*.
C. **Synonyms**: This group refers to instances in which the target terms *w*_1_ and *w*_2_ belong to the same UMLS concept. Each UMLS synonym set consists of a group of biomedical synonym concepts that express the same meaning. As shown in Figure 1, due to semantic ambiguity, biomedical concepts with several distinct meanings can be represented by several distinct synonym sets. For instance, *“Adenocarcinoma”* could have the same meaning as either *“Renal Cell Carcinoma”* (CUI C0007134) or *“Carcinoma in adenoma”* (CUI C0001418). Consequently, we consider these concepts as potential synonyms, which may or may not hold the same meaning depending on their context. To create the instances, we collect all the tuple pairs *t* = [(*s*_*i*_,*w*_*i*_),(*s*_*j*_,*w*_*j*_)] from *T* in which *w*_*i*_ and *w*_*j*_ both are present in a UMLS synonym set. We then assigned the label *True* to each instance if *w*_*i*_ and *w*_*j*_ are linked to the same UMLS CUI code, and *False* if they are not. Two examples of this group of instances are shown in Table 2.
D. **Label similarity**: Despite broad coverage of synonyms and semantic types, UMLS synonym sets still suffer a lack of a large number of reformed concepts that can be used in biomedical contexts. For instance, the concept “chronic pseudomonas aeruginosa infection” can be reformed as “chronic PA infection”, which is not covered by UMLS. To deal with this and to cover a wide range of target concepts with different formats in the dataset, we developed the fourth group of instances in which the corresponding terms have a high Levenshtein distance ratio (see examples 7 and 8 in Table 2). To create such instances, we retrieve all tuple pairs *t* = [(*s*_*i*_,*w*_*i*_),(*s*_*j*_,*w*_*j*_)] ∈ *T* in which the Levenshtein distance between *w*_*i*_ and *w*_*j*_ surpasses the threshold of 0.75. Each tuple *t* is marked as *True* when *w*_*i*_ and *w*_*j*_ correspond to the identical UMLS entry, and *False* in the other case. The main idea behind this strategy was to include instances where target terms have similar surface forms but refer to different medical concepts. Two instances of this group are shown in Table 2. In example 7, both “piebald” and “piebaldism” refer to the same concept, whereas in example 8, “anemic” and “anaemia” refer to two different concepts.

## 3. Data Records

We divided the BioWiC instances into three main parts i.e., training set, development set, and test set, thereby establishing a structured and robust framework for model development and evaluation. To do so, we first built the test set including 2’000 instances with three constraints: 1) only one instance for each unique pair of target terms, 2) no sentence repetition between instances, and 3) no overlap between sentences and term pairs of the test set and training or development sets. The primary objective of rules 1 and 2 was to ensure a diverse range of term pairs and sentences in the test set. Rule 3 was also introduced to assess the generalization power of the language models, i.e., the model’s ability to adapt to new, previously unseen data. Taking into account the constraints mentioned, we randomly sampled a set of 2000 term pair instances from the groups defined in section 2.3.1 (800, 200, 800, and 200 samples for term identity, abbreviations, synonyms, and label similarity groups, respectively) to build the testing data set. Finally, we used the remaining instances to create the training set. General statistics of the different splits of BioWiC are reported in Table 3. In addition, following WiC, we balanced all the data splits in terms of the number of tags, i.e., 50% *True* and 50% *False*.

**Table 3.** General statistics of BioWiC, divided by splits. “term idn”, “abbs”, “syns”, and “label sim”, stand for “term identity”, “abbreviations”, “synonyms”, and “label similarity”, respectively. The number of words per instance is calculated using the BERT (bert-base-uncased) tokenizer.

| Split | Instance group |  |  |  |  | Words per sentence | Unique sentences | Unique term pairs | Unique CUIs | Unique semantic types | Unique semantic groups |
| --- | --- | --- | --- | --- | --- | --- | --- | --- | --- | --- | --- |
|  | Term idn | Abbs | Syns | Label sim | All |  |  |  |  |  |  |
| Train | 3606 | 2480 | 7072 | 3998 | 17156 | 22.8 | 14522 | 5243 | 4448 | 98 | 15 |
| Dev | 400 | 100 | 400 | 100 | 1000 | 22.8 | 1820 | 821 | 1052 | 81 | 14 |
| Test | 800 | 200 | 800 | 200 | 2000 | 23.0 | 4000 | 2000 | 1870 | 88 | 15 |
| All | 4806 | 2780 | 8272 | 4298 | 20156 | 22.9 | 20102 | 8064 | 5303 | 99 | 15 |

During the compilation of the training set, we adopted a simple approach where we only included examples of their corresponding sentences that did not exceed a certain frequency threshold. We built the training set with various thresholds, ranging from 1 to 200, to determine the most appropriate limit. As illustrated in Figure 3, the size of the training set, the number of unique concepts, and the number of semantic types in the training set varied based on these thresholds. It was observed that once a sentence recurrence surpassed 100 times, the incremental growth of the training set size as well as the number of unique concepts was marginal, registering below 2%. Furthermore, if the threshold is set higher, the number of unique semantic types included in the training set will not exceed 98. As a result, we chose 100 as our cut-off point.

**Figure 3.**
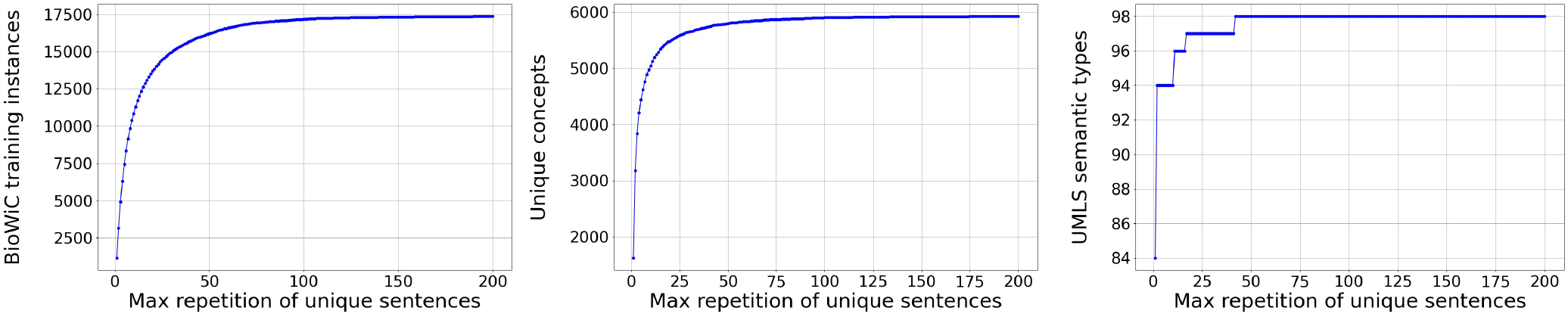
Impact of different thresholds for max sentence repetition in the training set. Left: Impact on the training set size; Center: Impact on the frequency of unique concepts; Right: Impact on the frequency of unique UMLS semantic types.

BioWiC dataset is available at https://github.com/hrouhizadeh/BioWiC. It comprises three distinct files: training set, development set, and test set, each of which is formatted in JSON. Each instance within the JSON files includes ten parts. The first two items, *term1* and *term2*, followed by *sentence1* and *sentence2*, correspond respectively to the two target terms and two sentences within each instance. The character-level positioning of target terms is defined by *start1* and *start2*, indicating the starting positions, and *end1* and *end2*, marking the end positions within their respective sentences. Furthermore, the *cat* attribute classifies each instance into one of the BioWiC groups, i.e., *term_identity, abbreviations, synonyms, or label_similairty*. Lastly, a binary *label* is attached to each instance, taking the value of either 1 (*True*) or 0 (*False*).

## 4. Technical Validation

### 4.1 Quality control

UMLS is known as a broadly used resource in the biomedical domain, covering a wide range of biomedical concepts. A key feature of UMLS is its capability to connect a wide range of concepts from different biomedical terminologies, such as SNOMED CT, LOINC, MeSH, RxNorm, etc. Through this mapping, one single code from a source terminology can be mapped to several UMLS CUI codes. For instance, MeSH code D020274, which represents “Depressive Disorder” is mapped to three distinct UMLS CUIs, C5671289, C0751871, and C0751872, for “Autoimmune Encephalitis”, “Autoimmune Diseases of the Nervous System” and “Immune Disorders, Nervous System”, respectively. In our dataset, there are instances where different CUI codes are assigned to the target concepts, resulting in the *False* label. However, the CUI codes and the confusion and same code in alternative ontologies, underlying concepts represented by those codes are equivalent. To prevent any confusion and to ensure the dataset’s reliability, we have employed a pruning strategy and removed the instances in which the target terms are mapped to multiple UMLS codes, while those UMLS codes correspond to the same code in another ontology. The process also involved eliminating any pairs whose CUIs are considered synonyms as per the *MRREL*.*RRF* file from UMLS. We also followed WiC and XL-WiC^20,22^ and filtered out all the pairs where one CUI is directly related to the other as a broader concept in the UMLS hierarchy.

### 4.2 Cross-mapping validation

To determine the quality of BioWiC, we extracted two random subsets of 100 instances (with 50 mutual instances) from the test set and asked two domain experts to label them. Both annotators were medical doctors with vast experience in semantic annotation. They were provided with a set of instructions including a short description of the task as well as a few examples of labeled instances. During the annotation process, no external information from UMLS or any other resources was provided to the experts. The annotators had Cohen’s Kappa score of 0.84 which is representative of the high quality of the dataset. An average human-level accuracy of 0.80 (0.80 and 0.81 for annotator 1 and annotator 2 respectively) was obtained through the annotation process, which can be viewed as the upper bound for model performance.

### 4.3 Dataset coverage

In this section, we focus on the scope of the dataset by studying the unique CUI codes and comparing them to the total CUI present in UMLS. Additionally, we investigate the semantic types within the dataset, examining both the number included and the proportions among them. Table 3 shows that BioWiC covers over 5,000 unique CUI codes from UMLS. Additionally, BioWiC includes almost 80% of UMLS semantic types, i.e., 99 out of 127, across different splits. This wide coverage is indicative of the dataset’s comprehensive and its potential as a valuable resource for biomedical research. In Figure 4, we present the ratio of the top 10 semantic types and semantic groups included in BioWiC. Additionally, Table 4 shows the frequency and proportion of target terms across different BioWiC splits, categorized by their token counts.

**Table 4.** Distribution of terms based on token count across different BioWiC splits, presented in counts and corresponding proportion. “4+” indicates terms with four or more tokens.

| Tokens per term | Train |  | Dev |  | Test |  | All |  |
| --- | --- | --- | --- | --- | --- | --- | --- | --- |
|  | # | % | # | % | # | % | # | % |
| 1 | 22011 | 64 | 1385 | 69 | 2987 | 74 | 26383 | 65 |
| 2 | 8893 | 26 | 458 | 23 | 784 | 20 | 10135 | 25 |
| 3 | 2406 | 7 | 118 | 06 | 166 | 4 | 2690 | 7 |
| 4+ | 1030 | 3 | 39 | 02 | 63 | 2 | 1132 | 3 |
| All | 34340 | 100 | 2000 | 100 | 4000 | 100 | 40340 | 100 |

**Figure 4.**
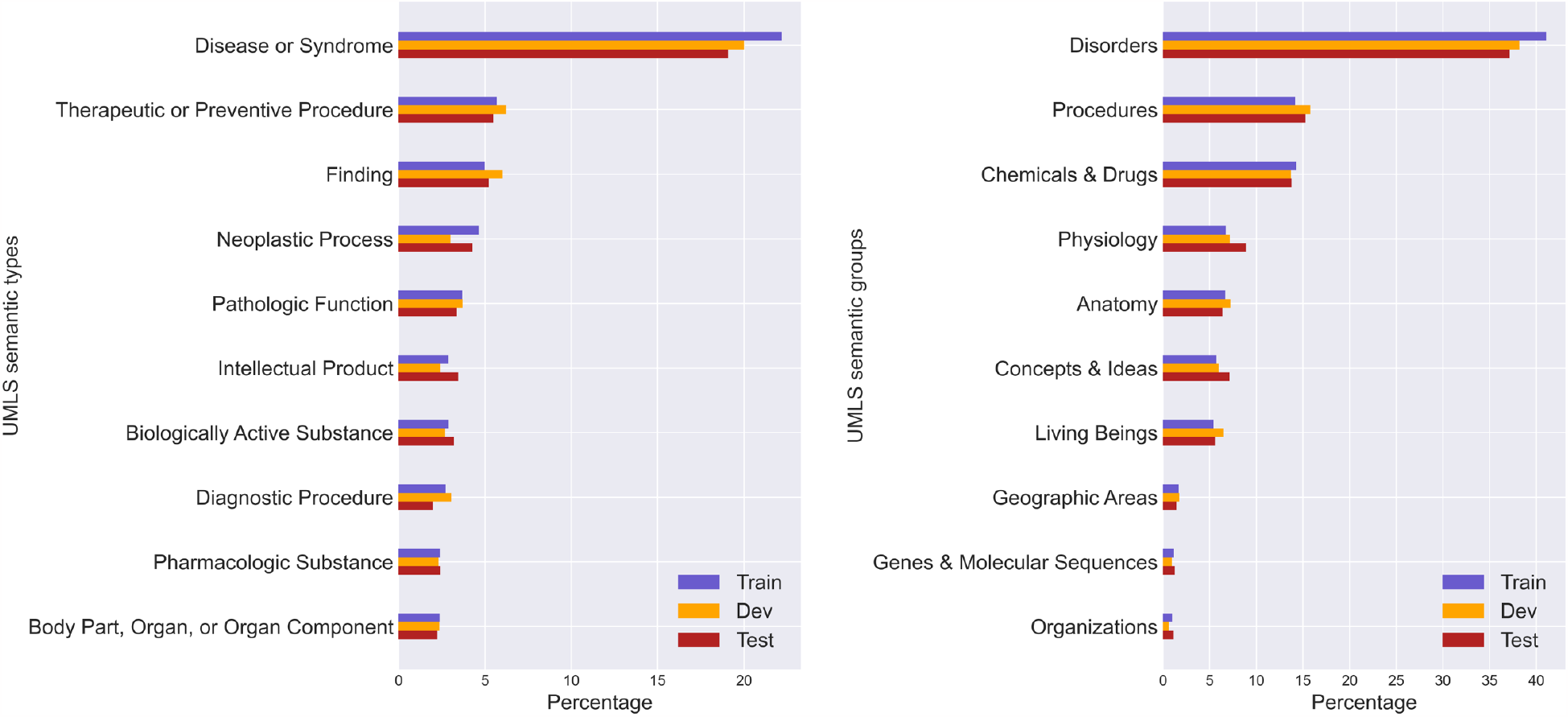
Distribution of UMLS semantic types and semantic groups in BioWiC. Left: Top 10 semantic types; Right: Top 10 semantic groups.

### 4.4 Baseline experiments

We have implemented several baseline models, covering all the SuperGLUE^21^ benchmark suites. Considering that all divisions of BioWiC are balanced in terms of positive and negative instances, we take the same approach as WiC^20^ and use the *accuracy* metric to measure the performance of different models. This is determined by the percentage of correctly predicted cases (whether they are true positives or true negatives) compared to the total number of samples. The baselines include:

#### Random

We provide a lower bound for the performance by randomly assigning a class to each instance.

#### GloVe

In this baseline, we used GloVe-840B^34^ pre-trained embeddings. We averaged token embeddings to represent each sentence and fed the resulting feature vector to an MLP classifier (with 128 neurons in the hidden layer and one neuron in the output layer).

#### Bi-LSTM

We also trained a BiLSTM model (with 128 hidden units) to capture both the forward and backward context information of the sentence. The BiLSTM model output was fed into a fully connected layer with one output neuron for binary classification.

#### BERT

We explored the performance of several BERT-based models to provide stronger baselines for the BioWiC task. To evaluate well language model’s performance generalized to concepts of the biomedical domain, our baseline includes three general transformer-based language models – BERT^35^, RoBERTa^36^, and ELECTRA^37^. In addition, to assess the effect of prior knowledge of language models on biomedical concept representation, we evaluated the performance of three language models pre-trained with biomedical and clinical data – BioBERT^38^, Bio_ClinicalBERT^39^, and SciBERT^40^ trained on PubMed abstracts and PubMed Central, the MIMIC-III database^41^, and papers from Semantic Scholar (mostly in the biomedical domain), respectively. To fine-tune each model, we used the Sentence-BERT^42^ framework, which incorporates siamese and triplet network architectures to generate semantically meaningful embeddings. We pre-processed each input sentence by enclosing the target terms within double quotes, emphasizing their significance, and fed the modified sentences into the BERT architecture for further processing.

#### Llama-2

We also conducted experiments using two different versions of the Llama-2 language model, i.e., Llama-2-7b and Llama-2-13b^43^. Our experiments involve a few-shot approach where the language model receives a small number of examples before making predictions and a fine-tuning approach, where we utilized the BioWiC instances to fine-tune the language models.

#### BERT/Llama-2 ++

We conducted additional experiments where we incorporated the general domain data from the WiC dataset^20^ as additional training data for fine-tuning the transformer-base models. By expanding our training data with extra instances from the general domain, we aim to explore the potential benefits of leveraging diverse sources of information for the BioWiC task.

### 4.5 Results

The performance of the baseline models on the BioWiC benchmark is presented in Figure 5. The results indicate that the state-of-the-art language models fine-tuned on the BioWiC training set, surpass the random baseline by a margin of 18% to 26% (*p*-value < 0.001). Both GloVe and BiLSTM baselines are unable to compete with the fine-tuned large language models. Overall, Llama-2-70b outperforms all competing methods, achieving the highest accuracy. The closest to the Llama-2-70b model in terms of accuracy are BioBERT, BioBERT ++, and SciBERT ++, which Llama-2-70b outperforms by 2% (*p*-value = 0.04). It is worth noting that in contrast to the different variations of the Lamma-2 language model, which are pre-trained on general domain corpora, BioBERT is pre-trained on large biomedical data, allowing it to understand complex biomedical texts effectively^38^. However, Llama-2-70b achieves state-of-the-art performance, illustrating its high capability for adapting to the task of representing biomedical terms in context.

**Figure 5.**
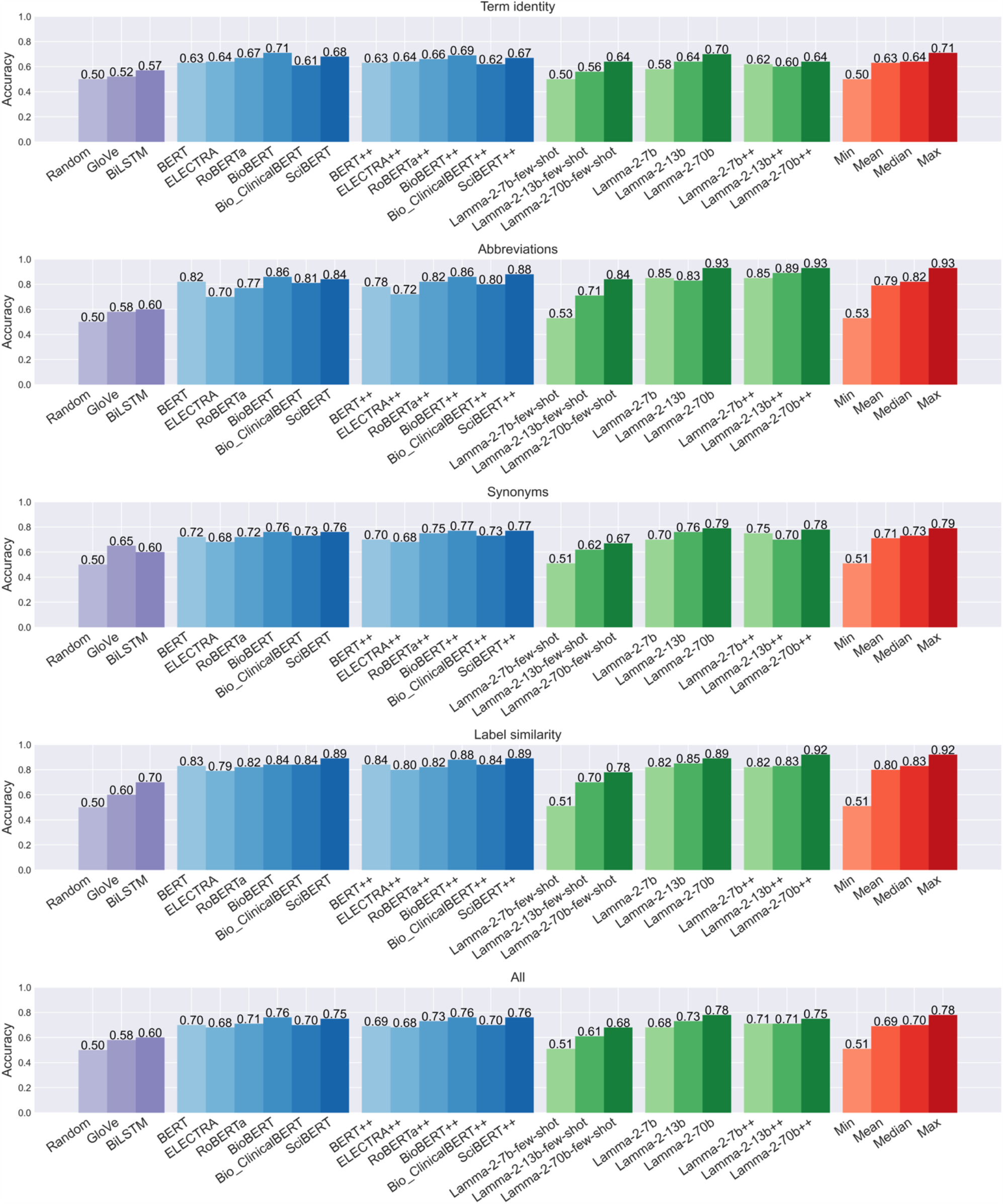
Accuracy of the baseline models on the BioWiC test set. ++ indicates that data from WiC was added to the training set. Min, mean, median, and max statistics exclude the random performance.

In our analysis of different Llama-2 models, we observe a significant difference in performance depending on the method used in our evaluation, i.e., few-shot learning or fine-tuning. As shown in Figure 5, Llama-2-7b surpassed the random baseline by a slight margin in the few-shot setting; however, its performance increased by 17% upon fine-tuning (*p*-value < 0.001). This pattern of performance boost was consistent with the other Llama-2 variants. Specifically, after the fine-tuning process, the accuracy of Llama-2-13b improved from 0.61 to 0.73 (*p*-value < 0.001), while Llama-2-70b experienced an increase from 0.68 to 0.78 (*p*-value < 0.001). These observations emphasize the crucial role of the fine-tuning phase in enhancing the contextualized representation of biomedical terms. Additionally, our results are consistent with a prior study^44^, which demonstrated that the GPT-3 language model failed to surpass random baseline performance on the WiC dataset under a few-shot evaluation.

Comparing the performance of different BERT-based models shows that BioBERT and SciBERT achieve the highest performance among different groups of the test set. Overall, BioBERT outperforms SciBERT by a slight margin of 1% accuracy, i.e., 0.75 and 0.76 (*p*-value = 0.04), respectively. The potential reason for the superior performance of BioBERT and SciBERT can be attributed to their pre-training phase on large biomedical corpora. This provides them with an in-depth knowledge of biomedical terminologies and concepts, leading to more accurate representations of terms and expressions when compared to BERT-based models pre-trained on the general domain corpora^38^. Surprisingly, Bio_ClinicalBERT performance is similar to the general domain BERT models and does not align with other superior biomedical BERT variants.

Further analysis of the results for different groups indicates that the “term identity” and “synonyms” groups present a greater challenge compared to the other groups for all models. Regarding model performance for the “label similarity” group, it is plausible that minor changes in term structure carry meaningful distinctions in biomedical contexts. Models might utilize structural alterations, such as the addition of suffixes or prefixes, influencing the meanings of terms. This understanding of term structure can be particularly relevant and beneficial for performance in the “label similarity” group. As for the “abbreviations” group, it is important to note that abbreviations are commonly used in the biomedical domain. The models may have come across these abbreviations (along with their full form) in various contexts during both the pre-training and fine-tuning phases. This exposure to abbreviations in diverse settings helps the models to effectively learn and capture their meanings. The group of “synonym” instances appears to be more difficult for models to handle. This might be because, in the biomedical field, a single term can have multiple synonyms with varied forms and each synonym can have multiple meanings (as shown in Figure 1) which makes it hard for the models to recognize synonym terms with different shapes across different contexts. For the “term identity” group, since this group of instances doesn’t present any difference between the target terms, the models cannot rely on lexical cues and must prioritize the comprehension of the contextual information from the surrounding context, which makes the task more challenging.

In our study, we also conducted experiments in which we incorporated general domain training data from WiC^20^ into our dataset (denoted by adding ++ to the name of the language model). We observe slight fluctuations in the performance of the models when merging general and biomedical domain datasets. It could be possibly explained by the fact that the model faces potential distribution shifts due to the distinct nature of each domain. Despite the increased volume of training data, this misalignment in data distributions can offset the advantages of the added samples. Thus, while the combined dataset is larger, it may not necessarily lead to improved model performance in the biomedical context.

#### 4.5.1 Alternative evaluation scenarios

To gain a deeper understanding of how models perform in the BioWiC benchmark, we analyzed their performance in two alternative scenarios. First, we assessed how the data distribution impact their results. Here, we considered seen and unseen data distributions. Second, we assessed what is the influence of the training corpus on the performance. Differently, in this scenario, we are interested to see whether learning from general corpus examples would enable models to generalise to the biomedical domain.

##### Seen vs unseen

In this analysis, the aim is to evaluate the variation in performance based on whether the target terms in the instances have been previously seen during training or not. For this purpose, we used the models fine-tuned on the BioWiC training set and divided the test set into two categories: “seen” and “unseen”. The first category includes instances where the model has been exposed to at least one of the target terms during training, while the second category involves instances where both target terms are new to the model. Table 5 reports the number and proportion of seen and unseen data across different groups within the BioWiC test set. Note that term pairs (the two target terms of each instance) and the sentences in the test set are unique and were not presented to the model during its training phase.

**Table 5.**
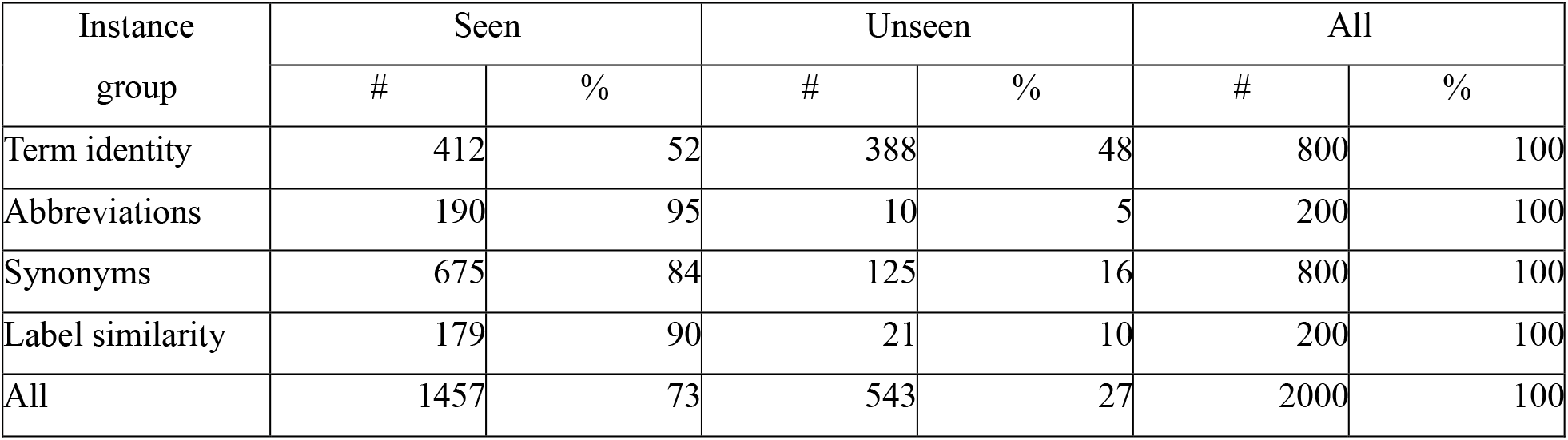
Distribution of seen and unseen instances in different groups of BioWiC test set.

| Instance group | Seen |  | Unseen |  | All |  |
| --- | --- | --- | --- | --- | --- | --- |
|  | # | % | # | % | # | % |
| Term identity | 412 | 52 | 388 | 48 | 800 | 100 |
| Abbreviations | 190 | 95 | 10 | 5 | 200 | 100 |
| Synonyms | 675 | 84 | 125 | 16 | 800 | 100 |
| Label similarity | 179 | 90 | 21 | 10 | 200 | 100 |
| All | 1457 | 73 | 543 | 27 | 2000 | 100 |

Table 6 shows the accuracy of different models, fine-tuned on the BioWiC training set when tested on seen and unseen data sets. As we can see, the models exhibit a significant decline in performance, i.e., between 5% and 13%, when classifying unseen instances. Interestingly, models demonstrate improved performance on the unseen data in the “abbreviation” groups, aligning with the notion that abbreviations are prevalent across contexts and models may possess prior knowledge in this aspect. Overal, the findings suggest that there is huge scope for improvement in this field, particularly as the performance of models decreases when encountering novel data.

**Table 6.** Comparative analysis of model accuracy on BioWiC test set. Left: performance of the models trained using BioWiC on the seen data vs unseen data distributions. Right: performance using WiC or BioWiC as training set.

| Model | Distribution |  | Training set |  |
| --- | --- | --- | --- | --- |
|  | Seen | Unseen | WiC | BioWiC |
| BERT | 0.72 | 0.67 | 0.63 | 0.70 |
| ELECTRA | 0.69 | 0.64 | 0.62 | 0.68 |
| RoBERTa | 0.74 | 0.64 | 0.63 | 0.71 |
| BioBERT | 0.79 | 0.67 | 0.64 | 0.76 |
| Bio_ClinicalBERT | 0.73 | 0.62 | 0.62 | 0.70 |
| SciBERT | 0.77 | 0.68 | 0.64 | 0.75 |
| Llama-2-7b-fine-tuned | 0.72 | 0.59 | 0.50 | 0.68 |
| Llama-2-13b-fine-tuned | 0.76 | 0.64 | 0.51 | 0.73 |
| Llama-2-70b-fine-tuned | 0.80 | 0.70 | 0.51 | 0.78 |

##### Cross-domain analysis

We conducted additional experiments to assess the performance of language models when fine-tuned exclusively on data from the general domain, specifically WiC. The results indicate that all models experience a substantial decrease in performance when fine-tuned only with WiC data (Table 6). This highlights the importance of the training data provided by BioWiC in enhancing the ability of language models in the representation of different forms of concepts within the biomedical field. Furthermore, this suggests that the differences in terminology and linguistic patterns between the biomedical and general domains might be a reason why models fine-tuned on BioWiC exhibit superior performance.

## 5. Usage Notes

The primary objective of this study is to develop a novel biomedical dataset, BioWiC, introducing unique challenges for biomedical concept representation. Using three manually annotated biomedical entity linking datasets, we developed four groups of instances i.e., term identity, abbreviations, synonyms, and label similarity, to present different challenges in the textual biomedical data to the models. We conducted several experiments using the BioWiC dataset to both train and evaluate state-of-the-art discriminative and generative large language models. These experiments provide robust baselines for future research in this field. We suggest that BioWiC provides a valuable resource for natural language processing research in the biomedical domain by enabling the evaluation and development of algorithms for biomedical concept representation.

A key attribute of BioWiC is its flexibility and scalability. Unlike WSD and entity linking that is restricted to concepts covered by existing knowledge graphs, BioWiC can be expanded independently of such resources. This is because expanding the dataset for a novel concept can be accomplished by annotating instances where two sentences contain the target concept, regardless of whether or not, it is included in any existing knowledge graph resource. This flexibility allows for continual evolution and improvement, independent of updates to standardized resources, providing a more comprehensive and up-to-date resource for research in the biomedical field.

The proposed benchmark has certain limitations that should be taken into consideration. The breadth of coverage of concepts is rather limited as BioWiC only deals with a small subset of the concepts present in the biomedical domain, i.e., 5’000 CUIs out of 4.5M CUI codes available in UMLS. Moreover, it may not be adequate for certain use cases that require a specific coverage of concepts, e.g., genomics and proteomics. Additionally, our benchmark is currently designed to work with medical documents written in English only. Lastly, it is a static benchmark, in the sense that it does not currently provide a seamless platform (i.e., web service) for users to contribute to it through crowd-sourcing. This limits the ability to keep the benchmark up-to-date and reflective of the latest developments in the biomedical domain. These limitations can be addressed in future versions of the benchmark.

## 6. Code availability

The entire process, including the development of the dataset and the conduction of experiments, was implemented using the Python programming language. The complete code and dataset are hosted on GitHub at: https://github.com/hrouhizadeh/BioWiC. In addition, the dataset is available at the Hugging Face repository at: https://huggingface.co/datasets/hrouhizadeh/BioWiC.

## 7. Conclusion

This paper proposes BioWiC, a novel benchmark for evaluating contextualized word embeddings across the biomedical domain. The benchmark is formulated as a binary classification task, in which the objective is to determine the semantic equivalence of terms in distinct biomedical contexts. Using three manually annotated datasets, we developed four groups of term-in-context instances to cover different semantic and concept representation challenges in the biomedical domain. The inter-annotator agreement based on a subset of the BioWiC test set (0.84) indicates that it can be used as a high-quality benchmark for training and evaluating biomedical concept representation models. We also evaluated a variety of general and domain-specific state-of-the-art language models on the benchmark and showed that discriminative and generative models achieve similar performance (0.76 vs. 0.78, respectively), despite a much larger number of parameters for the latter. Unlike WSD and entity linking, which are closely related tasks that deal with concept representation, the BioWiC task formulation does not rely on biomedical ontologies. This characteristic enables the benchmark to adapt to emerging biomedical concepts that are not yet included in existing knowledge organization systems. In the future, we plan to use different resources to extend BioWiC to different specific biomedical fields. Also, it would be interesting to develop and evaluate more complex models on the benchmark. Finally, we suggest that the task be applied to different languages, especially low-resource ones that are not covered by biomedical ontologies.

## Author contributions statement

H.R. and D.T. conceptualized the study, and H.R., A.Y, and B.Z. implemented the codes for the creation and evaluation of the dataset. J.E. and C.G. performed human annotation. The manuscript was drafted by H.R., D.T., and I.N., and edited by A.B, A.Y, and N.N. All authors reviewed the final version.

## References

1. Detroja, K., Bhensdadia, C. & Bhatt, B. S. A survey on relation extraction. Intell. Syst. with Appl. 200244 (2023).

2. Shi, J. et al. Knowledge-graph-enabled biomedical entity linking: a survey. World Wide Web 1–30 (2023).

3. French, E. & McInnes, B. T. An overview of biomedical entity linking throughout the years. J. Biomed. Informatic 104252 (2022).

4. Zhang, Y. et al. Neural network-based approaches for biomedical relation classification: A review. J. Biomed. Informatics 99, 103294, 10.1016/j.jbi.2019.103294 (2019).

5. Zhao, S., Su, C., Lu, Z. & Wang, F. Recent advances in biomedical literature mining. Briefings Bioinforma. 22, 10.1093/bib/bbaa057 (2020). Bbaa057, https://academic.oup.com/bib/article-pdf/22/3/bbaa057/37965584/bbaa057.pdf.

6. Magge, A. et al. DeepADEMiner: a deep learning pharmacovigilance pipeline for extraction and normalization of adverse drug event mentions on Twitter. J. Am. Med. Informatics Assoc. 28, 2184–2192, 10.1093/jamia/ocab114 (2021). https://academic.oup.com/jamia/article-pdf/28/10/2184/40408928/ocab114.pdf.

7. Song, B., Li, F., Liu, Y. & Zeng, X. Deep learning methods for biomedical named entity recognition: a survey and qualitative comparison. Briefings Bioinforma. 22, 10.1093/bib/bbab282 (2021). Bbab282, https://academic.oup.com/bib/article-pdf/22/6/bbab282/41089553/bbab282.pdf.

8. Donnelly, K. et al. Snomed-ct: The advanced terminology and coding system for ehealth. Stud. health technology informatics 121, 279 (2006).

9. Consortium, U. et al. Uniprot: the universal protein knowledgebase in 2021. Nucleic acids research 49, D480–D489(2021).

10. Erhardt, R. A., Schneider, R. & Blaschke, C. Status of text-mining techniques applied to biomedical text. Drug discoverytoday 11, 315–325 (2006).

11. Frénal, K., Kemp, L. E. & Soldati-Favre, D. Emerging roles for protein s-palmitoylation in toxoplasma biology. Int. J. forParasitol. 44, 121–131 (2014).

12. Alexopoulou, D. et al. Biomedical word sense disambiguation with ontologies and metadata: automation meets accuracy. BMC bioinformatics 10, 1–15 (2009).

13. Sung, M., Jeon, H., Lee, J. & Kang, J. Biomedical entity representations with synonym marginalization. arXiv preprintarXiv:2005.00239 (2020).

14. Navigli, R. Word sense disambiguation: A survey. ACM computing surveys (CSUR) 41, 1–69 (2009).

15. Moro, A., Raganato, A. & Navigli, R. Entity linking meets word sense disambiguation: a unified approach. Transactions Assoc. for Comput. Linguist. 2, 231–244 (2014).

16. Miftahutdinov, Z., Kadurin, A., Kudrin, R. & Tutubalina, E. Medical concept normalization in clinical trials with drug anddisease representation learning. Bioinformatics 37, 3856–3864 (2021).

17. Tutubalina, E., Miftahutdinov, Z., Nikolenko, S. & Malykh, V. Medical concept normalization in social media posts with recurrent neural networks. J. biomedical informatics 84, 93–102 (2018).

18. Niu, J., Yang, Y., Zhang, S., Sun, Z. & Zhang, W. Multi-task character-level attentional networks for medical concept normalization. Neural Process. Lett. 49, 1239–1256 (2019).

19. Limsopatham, N. & Collier, N. Normalising medical concepts in social media texts by learning semantic representation. In Proceedings of the 54th annual meeting of the association for computational linguistics (volume 1: long papers),1014–1023 (2016).

20. Pilehvar, M. T. & Camacho-Collados, J. WiC: the word-in-context dataset for evaluating context-sensitive meaning representations. In Proceedings of the 2019 Conference of the North American Chapter of the Association for Computational Linguistics: Human Language Technologies, Volume 1 (Long and Short Papers), 1267–1273, 10.18653/v1/N19-1128 (Association for Computational Linguistics, Minneapolis, Minnesota, 2019).

21. Wang, A. et al. Superglue: A stickier benchmark for general-purpose language understanding systems. Adv. neural information processing systems 32 (2019).

22. Raganato, A., Pasini, T., Camacho-Collados, J. & Pilehvar, M. T. XL-WiC: A multilingual benchmark for evaluating semantic contextualization. In Proceedings of the 2020 Conference on Empirical Methods in Natural Language Processing (EMNLP), 7193–7206, 10.18653/v1/2020.emnlp-main.584 (Association for Computational Linguistics, Online, 2020).

23. Loureiro, D. et al. Tempowic: An evaluation benchmark for detecting meaning shift in social media. arXiv preprint arXiv:2209.07216 (2022).

24. Miftahutdinov, Z. & Tutubalina, E. Deep neural models for medical concept normalization in user-generated texts. In Proceedings of the 57th Annual Meeting of the Association for Computational Linguistics: Student Research Workshop, 393–399, 10.18653/v1/P19-2055 (Association for Computational Linguistics, Florence, Italy, 2019).

25. Liu, F., Shareghi, E., Meng, Z., Basaldella, M. & Collier, N. Self-alignment pretraining for biomedical entity representations. arXiv preprint arXiv:2010.11784 (2020).

26. Angell, R., Monath, N., Mohan, S., Yadav, N. & McCallum, A. Clustering-based inference for biomedical entity linking. arXiv preprint arXiv:2010.11253 (2020).

27. Mohan, S. & Li, D. Medmentions: A large biomedical corpus annotated with umls concepts. In Automated Knowledge Base Construction (AKBC) (2019).

28. Loureiro, D. & Jorge, A. M. Medlinker: Medical entity linking with neural representations and dictionary matching. In European Conference on Information Retrieval, 230–237 (Springer, 2020).

29. Mohan, S., Angell, R., Monath, N. & McCallum, A. Low resource recognition and linking of biomedical concepts from a large ontology. In Proceedings of the 12th ACM Conference on Bioinformatics, Computational Biology, and Health Informatics, 1–10 (2021).

30. Li, J. et al. Biocreative v cdr task corpus: a resource for chemical disease relation extraction. Database 2016 (2016).

31. Dogan, R. I., Leaman, R. & Lu, Z. Ncbi disease corpus: a resource for disease name recognition and concept normalization. J. biomedical informatics 47, 1–10 (2014).

32. Sadvilkar, N. & Neumann, M. PySBD: Pragmatic sentence boundary disambiguation. In Proceedings of Second Workshop for NLP Open Source Software (NLP-OSS), 110–114, 10.18653/v1/2020.nlposs-1.15 (Association for Computational Linguistics, Online, 2020).

33. Stevenson, M. & Guo, Y. Disambiguation of ambiguous biomedical terms using examples generated from the umls metathesaurus. J. biomedical informatics 43, 762–773 (2010).

34. Pennington, J., Socher, R. & Manning, C. D. Glove: Global vectors for word representation. In Proceedings of the 2014 conference on empirical methods in natural language processing (EMNLP), 1532–1543 (2014).

35. Devlin, J., Chang, M.-W., Lee, K. & Toutanova, K. Bert: Pre-training of deep bidirectional transformers for language understanding. arXiv preprint arXiv:1810.04805 (2018).

36. Liu, Y. et al. Roberta: A robustly optimized bert pretraining approach. arXiv preprint arXiv:1907.11692 (2019).

37. Clark, K., Luong, M.-T., Le, Q. V. & Manning, C. D. Electra: Pre-training text encoders as discriminators rather than generators. arXiv preprint arXiv:2003.10555 (2020).

38. Lee, J. et al. Biobert: a pre-trained biomedical language representation model for biomedical text mining. Bioinformatics 36, 1234–1240 (2020).

39. Alsentzer, E. et al. Publicly available clinical BERT embeddings. In Proceedings of the 2nd Clinical Natural Language Processing Workshop, 72–78, 10.18653/v1/W19-1909 (Association for Computational Linguistics, Minneapolis, Minnesota, USA, 2019).

40. Beltagy, I., Lo, K. & Cohan, A. Scibert: A pretrained language model for scientific text. arXiv preprint arXiv:1903.10676 (2019).

41. Johnson, A. E. et al. Mimic-iii, a freely accessible critical care database. Sci. data 3, 1–9 (2016).

42. Reimers, N. & Gurevych, I. Sentence-bert: Sentence embeddings using siamese bert-networks. arXivpreprint arXiv:1908.10084 (2019).

43. Hugo Touvron, et al. 2023. Llama 2: Open foundation and fine-tuned chat models. arXiv preprint arXiv:2307.09288 (2023)

44. Brwon T. et al. Language models are few-shot learners. Adv. neural information processing systems 33, 1877–1901 454 (2020)

